# Independent Trafficking and Diverse Stability of XyG-Synthesizing Glycosyltransferases in Distinct Golgi cisternae

**DOI:** 10.64898/2026.04.15.718703

**Authors:** Ning Zhang, Kayla Uthe, Olga A Zabotina

## Abstract

XyG-synthesizing glycosyltransferases (GTs) are localized in Golgi, and their protein-protein interactions suggest the formation of multiprotein complexes; however, the mechanisms underlying protein complex assembly and transportation, protein stability and degradation remain unknown. We have uncovered that protein-protein interactions among XyG-synthesizing GTs are not prerequisites for Golgi localization and different GTs are first delivered to Golgi as independent proteins. By employing the transient expression of YFP-fused GTs along with a cis-Golgi marker and generating fluorescence intensity profiles, we demonstrated the differential distribution of GTs in Golgi apparatus. We treated *Arabidopsis* seedlings expressing GTs with cycloheximide (CHX) to estimate half-lives, The GTs exhibit distinct half-lives, and based on their turnover rates, divided into two groups. Our findings revealed that cellulose synthase-like C4 (CSLC4), galactosyltransferase (MUR3), and fucosyltransferase (FUT1) exhibit longer stability. In contrast, XyG xylosyltransferases XXT1, XXT2, XXT5, and galactosyltransferase XLT2 are significantly shorter-living proteins. XyG-synthesizing proteins traffic independently to Golgi, exhibiting distinct sub-Golgi localization that shapes multiprotein complex assembly in specific cisternae. This spatial organization governs partner GT access and residence time for functional efficiency, while GT half-life variations regulate stability and interaction dynamics. Collectively, these factors provide a critical framework for independent operation and coordinated organization of Golgi-resident XyG-synthesizing proteins.

**Highlight:** Glycosyltransferase traffic independently to the Golgi, exhibit distinct sub-Golgi localization and variable half-lives that reveal the mechanism of protein interactions, stability, and assembly of multiprotein complexes for stepwise synthesis.

## Introduction

Xyloglucans (XyGs) are highly branched polysaccharides and are the main hemicellulosic component in the primary walls of flowering plants (Scheller and Ulvskov, 2010). Various types of XyG structures are found in different plant species and tissues. In *Arabidopsis*, the biosynthesis of XLFG-type XyGs has been studied most intensively (Pauly and Keegstra, 2016, Preprint; Julian and Zabotina, 2022). The cellulose synthase-like C4/5/6/8/12 (CSLC4/5/6/8/12) proteins catalyze the synthesis of the β-1,4-linked xyloglucan glucan backbone (Cocuron et al., 2007; Kim et al., 2020). Two xyloglucan xylosyltransferases, XXT1 and XXT2, add the xylosyl residues on the first and second glucose (Cavalier et al., 2008; Zabotina et al., 2012), whereas XXT3, XXT4, and XXT5 specifically add the xylosyl residues on the third glucose in the XyG glucan backbone (Zabotina et al., 2009; Vuttipongchaikij et al., 2012; Zhang et al., 2023). Galactosyltransferases XLT2 (Jensen et al., 2012) and MUR3 (Madson et al., 2003; Peña et al., 2004; Tamura et al., 2005; Kong et al., 2015) add galactoses to the second and third xyloses to extend the side chains of XyGs. Fucosyltransferase FUT1 further extends the side chain on the third glucose, finalizing the F-chain in XLFG-type XyGs (Vanzin et al., 2002; Peña et al., 2004).

The biosynthesis of the highly branched polysaccharides and complex N- and O-glycans in various glycoconjugates occurs in the Golgi. This process requires the cooperation of multiple GTs, which function as homodimers, heterodimers, or larger complexes (Anderson, 2016; Strasser et al., 2021; Julian and Zabotina, 2022). Moreover, some GTs initiate assembly into homodimers, heterodimers, or higher-order complexes in the ER, followed by their trafficking to the Golgi apparatus as multiple protein complexes to execute their core enzymatic functions (Jiang et al., 2016; Zeng et al., 2016). In the case of xylan biosynthesis in *Asparagus officinalis,* the collaboration of three xylosyltransferases, namely IRREGULAR XYLEM9 (AoIRX9), AoIRX14A, and AoIRX10 has been shown. These enzymes have been demonstrated to engage in protein-protein interactions that play a pivotal role in their ER-Golgi exportation, where AoIRX9 drives this complex to the Golgi apparatus. (Zeng et al., 2016). Similarly, in wheat (*Triticum aestivum*), the putative xylan glycosyltransferase TaGT43-4 interacts with the glycosyltransferase TaGT47-13, mutase TaGT75-3, mutase TaGT75-4, and protein TaVER2, collectively forming a heterocomplex within the ER (Jiang *et al*., 2016). TaGT43-4 recruits this heterocomplex to the trans-Golgi, presumably functioning as a scaffold protein (Jiang et al., 2016). Protein-protein interactions have also been demonstrated among the GTs involved in XyG biosynthesis (Chou et al., 2012, 2015; Lund et al., 2015). The integral membrane protein CSLC4 forms heterodimers with most type II membrane GTs involved in synthesizing the XyG side chains: XXTs, XLT2, MUR3, and FUT1 (Chou et al., 2012, 2015; Lund et al., 2015). These findings strongly suggest that multiprotein complexes of XyG-synthesizing GTs, with the unknown mechanism of protein and protein complex transportation, composition in the Golgi and stoichiometry.

Knowledge of protein stability offers clues to understanding protein turnover and the timing of their function. Studies targeting the half-life of GTs are limited and have been performed mostly in human or yeast cells. Although scarce, these investigations have revealed significant variability in the half-life of GTs, ranging from minutes to hours. Cycloheximide (CHX) interferes with protein synthesis in ribosomes, thereby inhibiting the cell’s total protein synthesis. This inhibitor is frequently utilized to study the half-life and turnover of diverse proteins. For example, ST3 β-galactoside α-2,3-sialyltransferase 4 (ST3GAL4), which participates in the α-2,3-sialylation of N-glycans (Gillespiesq et al., 1992), shows high stability and does not change significantly after 36 h of CHX treatment (Kitano et al., 2021). Golgi-localized β-1,6-N-acetylglucosaminyltransferase 3 (GCNT3), which catalyzes the attachment of GlcNAc to branch and extend the various O-glycans (Li et al., 2018; Gupta et al., 2020; Zhao et al., 2022), was detected by western blot after 36 h of exposure to CHX (Sumardika et al., 2018).

The level of Growth Regulation by Estrogen in the Breast Cancer 1 (GREB1) protein, a putative glycosyltransferase, exhibits a short half-life of ∼6 h following CHX treatment and is barely detectable after 12 h (Myoung Shin et al., 2021). Among human Multiple UDP-glucuronosyltransferases (UGTs) 2 subfamily proteins, UGT2B4, UGT2B7, and UGT2B15 are long-lived, maintaining levels after 12 h CHX exposure (Barbier et al.,1999, 2000; Turgeon et al., 2001), whereas the fourth GT, UGT2B17, is short-lived and is undetectable after 12 h of CHX treatment of cells (Turgeon et al., 2001). On the other hand, the short-term protein stability of UGT2B20 has been reported, with a 50% loss of enzymatic activity observed after 30 min of CHX treatment (Barbier et al., 1999, 2000). O-GlcNAc transferases (OGTs) stability varies by cell type, with a ∼2 h half-life in HEK293 cells but ∼13 h in lymphoblastoid cells (Vaidyanathan et al., 2017; Peng et al., 2021). Similarly, ER-resident N-glycosylation heterooligomeric UDP-N-acetylglucosamine transferase Alg13p and Alg14p show stark differences, with unstable soluble Alg13p (<10 min half-life) comparable to stable membrane-bound Alg14p (30–60 min) (Averbeck et al., 2008). An investigation of Alg13p degradation via the proteasomes demonstrated that the interaction between Alg13p and Alg14p significantly increased the half-life of their complex, due to the membrane-bound Alg14p subunit stabilizing the complex (Averbeck et al., 2008). Collectively, these findings reveal diverse GT stabilities that influence multiprotein complex formation, longevity, and function, providing key insights into synthetic pathways that support cellular homeostasis and environmental responses.

To investigate the functional organization and protein stability of GTs in polysaccharide biosynthesis, we investigated the trafficking, Golgi distribution, and half-lives of seven *Arabidopsis* XyG-synthesizing GTs. First, we examined GT localization in single and multiple mutants of other XyG-biosynthetic GTs to determine whether protein–protein interactions are required for Golgi targeting. We then analyzed their spatial distribution within Golgi cisternae using fluorescence intensity profiles of YFP-GT fusions along with an mCherry-labeled cis-Golgi marker. Additionally, we investigated the stability of all XyG-synthesizing GTs by estimating their half-life using CHX. Together, the results from these experiments establish a framework for the assembly, organization, and dissociation of XyG-synthesizing multiprotein complexes.

## Materials and methods

### Plant growth conditions and plasmid construction

All T-DNA knockout (KO) mutant lines were obtained from ABRC (https://abrc.osu.edu): *xxt2* (Salk_101308) (Cavalier et al., 2008), *xxt1xxt2xxt5* (CS67828) (Zabotina et al., 2012), *cslc456812* (CS72438) (Kim et al., 2020), *xlt2* (GK-552C10-021687) (Jensen et al., 2012), *mur3-7* (SALK_127057) (Kong et al., 2015) and *fut1* (Salk_139678) (Jensen et al., 2012). Prof. Michael Hahn kindly provided the seeds of *xlt2*, *fut1*, and *mur3-7*, while Prof. Federica Brandizzi provided the seeds for the *cslc456812* quintuple mutant. To check the localization of GTs in the *Arabidopsis* Col-0 and mutant protoplasts, six-week-old plants grown in the growth chamber under the same conditions were used to prepare the protoplasts.

For the experiments with CHX treatment, the surface-sterilized Arabidopsis seeds were germinated and grown on sterile Petri dishes on 1% (w/v) agar with ½ MS media at pH 5.8 for 7 or 14 days in the growth chamber at 22°C and under long-day conditions (16 h:8 h, light: dark).

The genes XXT1 (AT3G62720), XXT2 (AT4G02500), XXT5 (AT1G74380), CSLC4 (AT3G28180), XLT2 (AT5G62220), MUR3 (AT2G20370), and FUT1 (AT2G03220) were amplified from previously reported plasmids (Chou et al., 2015) using gene-specific primers (Supplemental Table S1). The PCR products were digested with the corresponding restriction enzymes and inserted into expression vectors (pUBN-CFP, Grefen et al., 2010).

### Transient expression of XyG genes in *Arabidopsis* protoplasts

The pUBN::CFP and pUBN::YFP vectors (Grefen et al., 2010) were used to fuse CFP/YFP with the GT genes at their N-termini. The vectors pUBN::CFP-GTs and pUBN::YFP-GTs were purified from *E. coli* cells and used for transfections. About 20–30 leaves from the six-week-old Col-0 and various *Arabidopsis* mutants specified in the results section were collected for protoplast preparation. The purified vectors pUBN::CFP-GTs or pUBN::YFP-GTs and markers vector (Golgi-Y (YFP) / R (mCherry and ER-Y (YFP) / R (mCherry) (Nelson *et al*., 2007) were co-transformed into the protoplasts following the previously described protocols (Chou et al., 2012). After overnight incubation in the dark at room temperature, the fluorescence signals in the protoplasts were imaged using a confocal microscope (Leica SP5 system and Leica STED super-resolution, Leica Microsystems Inc, Deerfield, IL, USA) to confirm the localization of GTs.

### Localization analysis of the YFP fused GTs and the trans-Golgi marker ST-GFP with cis-Golgi marker cis-Golgi marker Golgi-R (mCherry)

pUBN::YFP-GTs vectors and trans-Golgi marker ST-GFP were transiently co-expressed with cis-Golgi marker Golgi-R (mCherry) (Nelson et al., 2007) in the protoplasts from Col-0 *Arabidopsis* and imaged using a confocal microscope (Leica SP5 system and Leica STED super-resolution, Leica Microsystems Inc, Deerfield, IL, USA). The plot profile intensity for YFP/GFP and mCherry fluorescence was analyzed by Image Fiji (Java 1.8.0_332 (64-bit)). The distance between the intensity peak of YFP/GFP and mCherry fluorescence was measured and statistically analyzed, representing the subcellular distribution and localization of GTs within the Golgi apparatus (Fig. 4).

### *Arabidopsis* stable transgenic lines

The recombinant vectors pUBN::CFP-GTs and pUBN::CFP were transformed into the corresponding single GT KO mutants and Col-0 via *Agrobacterium* using the floral dipping method (Clough and Bent, 1998). The transformants were selected on Petri dishes with hygromycin B, and the expression of GTs fused with CFP was confirmed by observing CFP fluorescence under a fluorescent microscope. The homozygous lines were selected and named CFP and CFP-GTs.

### GTs’ half-life determination

#### Analysis of the half-lives of GTs using western blot

To determine the protein contents using western blot, the fourteen-day-old *Arabidopsis* seedlings grown on the plates were collected and divided into eight samples of equal weight (1.5-2.5 g per sample). The *Arabidopsis* seedlings of each sample were submerged in 20 mL to 40 mL of ddH_2_O containing 500 µg/mL CHX and collected at the indicated time points. The treated seedlings were immediately ground with liquid nitrogen and suspended in 20 mL of the protein extraction buffer (50 mM HEPES, 3 mM DTT, 0.3 M sucrose, 65 mM NaCl, 3 mM EDTA, pH 7.5) containing 200 µL protease inhibitor cocktail (1 mM E-64, 1 mM Leupeptin, 100 mM AEBSF, 100 mM Benzamidine, and 100 mM PMSF in Methanol). The plant debris was filtered out using miracloth (Millipore Sigma, Lenexa, KS, USA, 4758551R), and the clean filtrates were transferred into 50 mL centrifuge tubes. The filtrates were centrifuged at 20,000 × g for 1 h at 4°C, and the supernatants were transferred to ultra-centrifuge tubes (Beckman Coulter Life Sciences, Indianapolis, IN, USA, Beckman #355618). After ultracentrifugation of the supernatants at 100,000 x g using the Ti70 rotor for 1 h at 4°C, the supernatants were discarded, and the pellets were resuspended in 1 mL of the protein extraction buffer. This step of washing the membrane pellets with the buffer was repeated three times. After washing the membrane pellet three times, it was resuspended in 30-50 µL of the protein extraction buffer containing 2% Triton X-100 and 1% NP-40 and shaken at 4°C overnight to solubilize membrane-bound proteins. The solubilized membrane proteins were transferred into 1.5 mL centrifuge tubes, placed on ice, and sonicated for further solubilization. After sonication, the tubes were centrifuged at 14,000 x g at 4°C, and the supernatants were collected. A Bio-Rad kit (Quick Start Bradford Dye reagent 1X; catalog, Thermo Fisher Scientific Inc, Waltham, MA, USA, no. 500-0205) was used to measure the total membrane protein content in the obtained supernatants, and an equal amount of total protein was used to prepare the samples for SDS-PAGE separation (10% Acrylamide). After SDS-PAGE, the proteins were transferred to nitrocellulose membranes (0.2 mm; Bio-Rad, Hercules, CA, USA) for immunodetection. Monoclonal anti-GFP antibodies (Covance, Princeton, NJ, USA) were used (1:6000 dilution) to detect the CFP-GT fusion proteins. The membranes were incubated with West Pico PLUS Chemiluminescent Substrate (Super Signal, Englewood, FL, USA) and visualized using ChemiDocXRS+ (Bio-Rad) system (Bio-Rad, Hercules, CA, USA). The intensities of the bands from the visualization were analyzed using Image Fiji.

### Analysis of the half-lives of GTs using fluorescent microscopy

Seven-day-old *Arabidopsis* seedlings were submerged in 500 µg/mL CHX and incubated for different periods. The intensity of the CFP was measured using a Zeiss Imager-upright microscope at consecutive time points. The root tips of *Arabidopsis* seedlings were forced on a 40× objective and held for the indicated exposure times. The background signal of Col-0 seedlings was measured and used as a negative control. Image Fiji was used to analyze the intensity of the CFP on all collected images. The three-square areas were randomly selected at each root tip, and the intensity of the CFP was estimated. The same imaging process was repeated at each time point for the CFP signals and the background of Col-0. The background signal value was subtracted from the intensity of the CFP at each time point image to remove the background contribution. The CFP intensities on the images from each time point were compared to the CFP intensity on the image at time zero, and the percentage of the signal was calculated.

### Statistical analysis

Data presented in Figure 3a-h and Figure 4a-h show the means and standard deviations (± SD) of three biological replicates of western blot analysis or fluorescent intensity (n=3). In Figure 3i and Figure 4i, the half-lives of GTs were estimated using results from each biological replication separately, and then statistically evaluated using ANOVA. The letter on the diagrams shown indicates the significant difference determined by one-way ANOVA (Tukey’s HSD p < 0.05). In Figure 2j, data are presented as the mean and standard deviation (± SD) of the distance between the intensity peak of YFP/GFP and mCherry (n=36), and the letter indicates the significant difference determined by one-way ANOVA (Tukey’s HSD p < 0.05).

## Results

### Localization of GTs expressed in mutant protoplasts

To investigate whether the XXT proteins are required for other XyG-synthesizing proteins to localize in the Golgi, the YFP-GT proteins were co-expressed with either Golgi-R (mCherry) or ER-R (mCherry) markers (Nelson et al., 2007) in protoplasts prepared from triple mutant *xxt1xxt2xxt5*. In the *xxt1xxt2xxt5* triple mutant, the CSCL4, XXTs (XXT1, XXT2, and XXT5), XLT2, MUR3, and FUT1 were localized in the Golgi, and their signal overlapped well with the Golgi marker but not the ER marker, as shown in merged images (Fig. 1).

**Figure 1.**
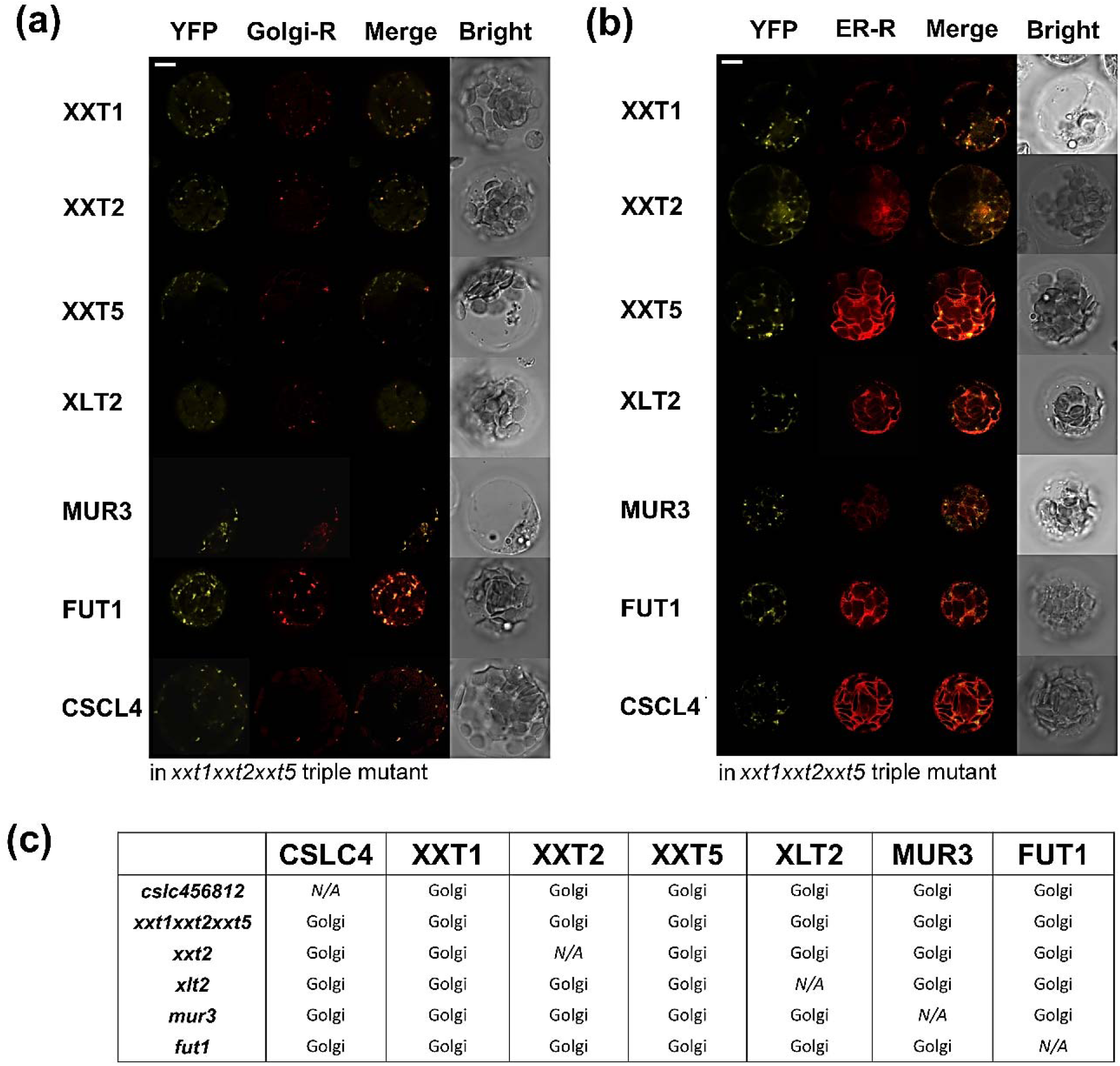
Transient co-expression of YFP-GTs with Golgi marker Golgi-R (mCherry) or ER-R (mCherry) in *Arabidopsis* protoplasts. (a) (b) The example of subcellular localization of YFP-GTs with Golgi marker Golgi-R (mCherry) or ER-R (mCherry) in *xxt1xxt2xxt5 Arabidopsis* protoplasts Bar, 10 μm. (c) The summary of the localization of other YFP-GTs in different KO mutant *Arabidopsis* protoplasts.

We further investigated the expression of all GTs in protoplasts prepared from various knockout (KO) mutant plants. In the protoplasts prepared from the *cslc456812* quintuple mutant plants, all proteins, XXT1, XXT2, XXT5, XLT2, MUR3, FUT1, and CSLC4 co-expressed with the Golgi or ER markers, showed localization in the Golgi (Fig. 1c; Supplemental Fig. S1). The CSLC family includes five members (CSLC4/5/6/8/12), shown to be involved in synthesizing the XyG glucan backbone and their different expression patterns indicate that they are functionally redundant but specific to tissue and developmental stages (Kim et al., 2020). Similar results were obtained for GTs expressed in protoplasts isolated from the *xlt2* mutant plants (Supplemental Fig. S2), as well as those from *mur3* (Supplemental Fig. S3), *fut1* (Supplemental Fig. S4) and *xxt2* (Supplemental Fig. S5) mutant plants: all seven proteins were localized in the Golgi, and none were found in the ER (Fig. 1; Supplemental Fig. S1-5). The expression pattern of each protein in the protoplasts prepared from the mutant plant of the same protein served as the positive control.

### Distribution of the XyG-synthesizing GTs within Golgi stacks

To further investigate the spatial distribution of the GTs in the Golgi apparatus, the localization of trans cisternae marker (ST-GFP) and YFP -fused GTs in Golgi stacks was examined in the Col-0 protoplasts expressing cis-Golgi marker (Golgi-R) (Nelson et al., 2007). The plot profile intensity for YFP/GFP and mCherry fluorescence was analyzed by Image Fiji. The distance between the peaks of YFP/GFP and mCherry signals was statistically analyzed. The results showed that XXT1 (Fig. 2g-i), XXT2 (Supplemental Fig. S6) and XXT5 (Supplemental Fig. S7) mainly remained in the cis/medial Golgi, CSLC4 (Supplemental Fig. S8), whereas XLT (Supplemental Fig. S9) exhibited the localization predominantly in the medial Golgi with some level of detection in cis- and trans-Golgi. In contrast, MUR3 (Supplemental Fig. S10) and FUT1 (Fig. 2e-f) predominantly localized in the medial/trans-Golgi (Fig. 2j).

**Figure 2.**
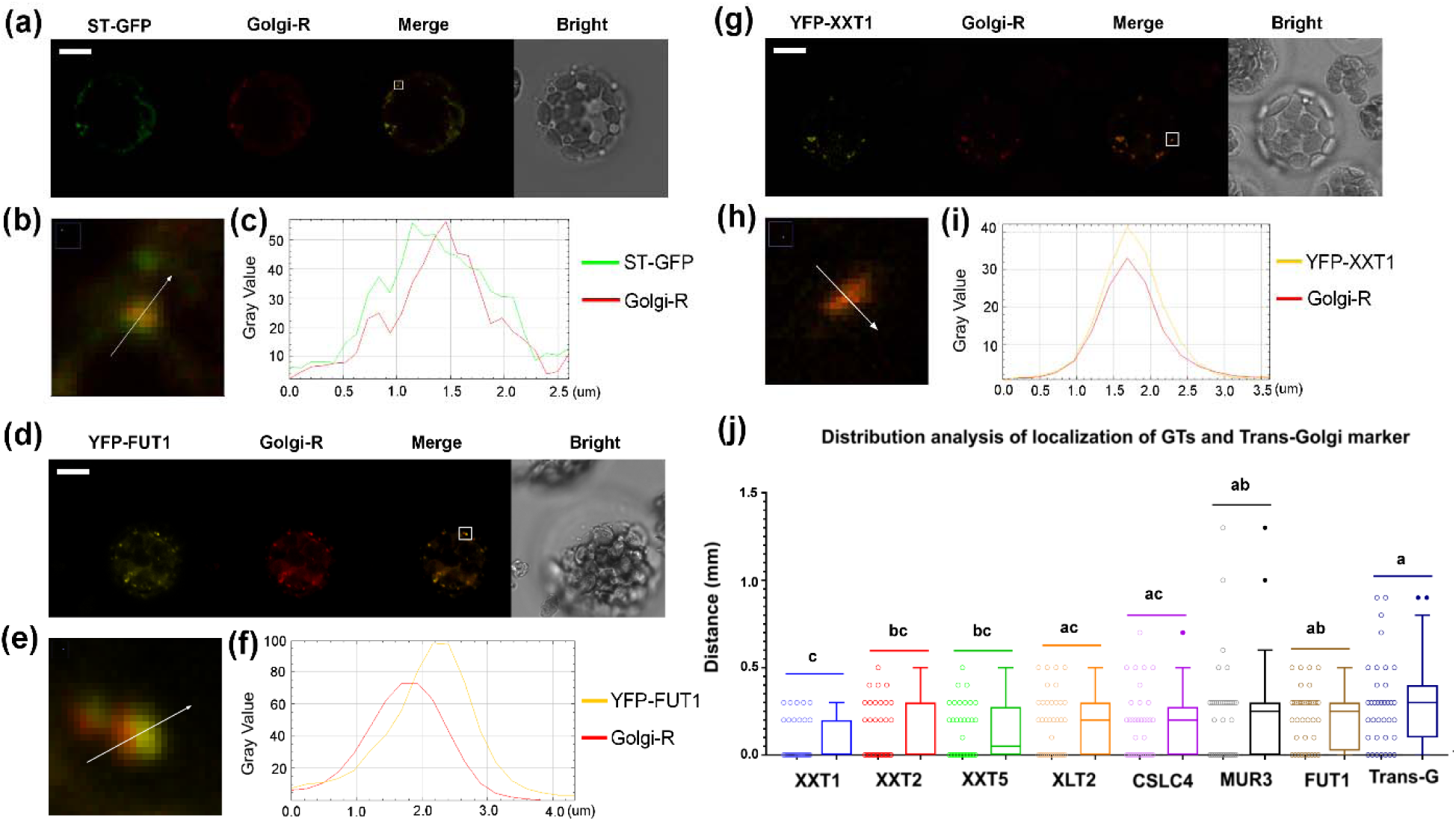
Distribution analysis of the localization of GTs and trans-Golgi marker ST-GFP. The YFP-GTs or trans-Golgi marker ST-GFP and cis-Golgi marker Golgi-R were co-expressed in the Col-0 protoplasts. Scale Bar, 10 μm. The plot profile intensity for YFP/GFP and mCherry fluorescence was analyzed by Image Fiji (n=36). The distance between the intensity peak of YFP/GFP and mCherry was analyzed. (a, d and g) The co-expression of trans-Golgi marker ST-GFP/YFP-XXT1/YFP-FUT1 and cis-Golgi marker Golgi-R in the Col-0 *Arabidopsis* protoplasts. The image in the white square was enlarged and shown in (b, e and h). (c, f and i) The plot profile intensity for GFP and mCherry fluorescence was analyzed by Image Fiji (n=36). The distance between the intensity peak of GFP and mCherry was analyzed. (j) The statistical analysis of distribution of the distance between intensity peak of YFP/GFP and mCherry (n=36) via one-way ANOVA. Letter indicates the significant difference by one-way ANOVA (Tukey’s HSD p < 0.05).

### Estimation of the half-lives of XyG-synthesizing GTs using Western blot

The inhibitor of protein synthesis, CHX, was utilized here to estimate the half-life of the XyG-synthesizing GTs. Total membrane protein fractions were prepared from fourteen-day-old *Arabidopsis* seedlings treated with CHX for different periods of time and analyzed by Western blot. The obtained results demonstrated significant variations in the half-lives of these proteins (Fig. 3). Based on the results, the XyG-synthesizing GTs can be divided into two distinct groups with significantly different stabilities: long-living and short-living GTs. The half-lives of XXT1 and XXT2 were about 30 mins, whereas the half-life of XXT5 was close to 25 mins, which is the shortest half-life for the GTs studied here (Fig. 3a, b and e). The half-life of the XLT2 protein was somewhat longer than that of the XXTs and was estimated to be about 50 mins (Fig. 3f). The long-living group of proteins included the CSLC4, MUR3, and FUT1 proteins, exhibiting several hours of half-life (Fig. 3c, d, and g). The half-life of CSLC4 was about 3.5h, and the half-life of FUT1 was about 4.5-5 h. MUR3 was estimated to have a half-life of approximately 5 h, the longest among XyG-synthesizing GTs. The half-life of CFP in Col-0 was used as a control and remained unchanged for at least 10 h after initiation of CHX treatment (Fig. 3h).

**Figure 3.**
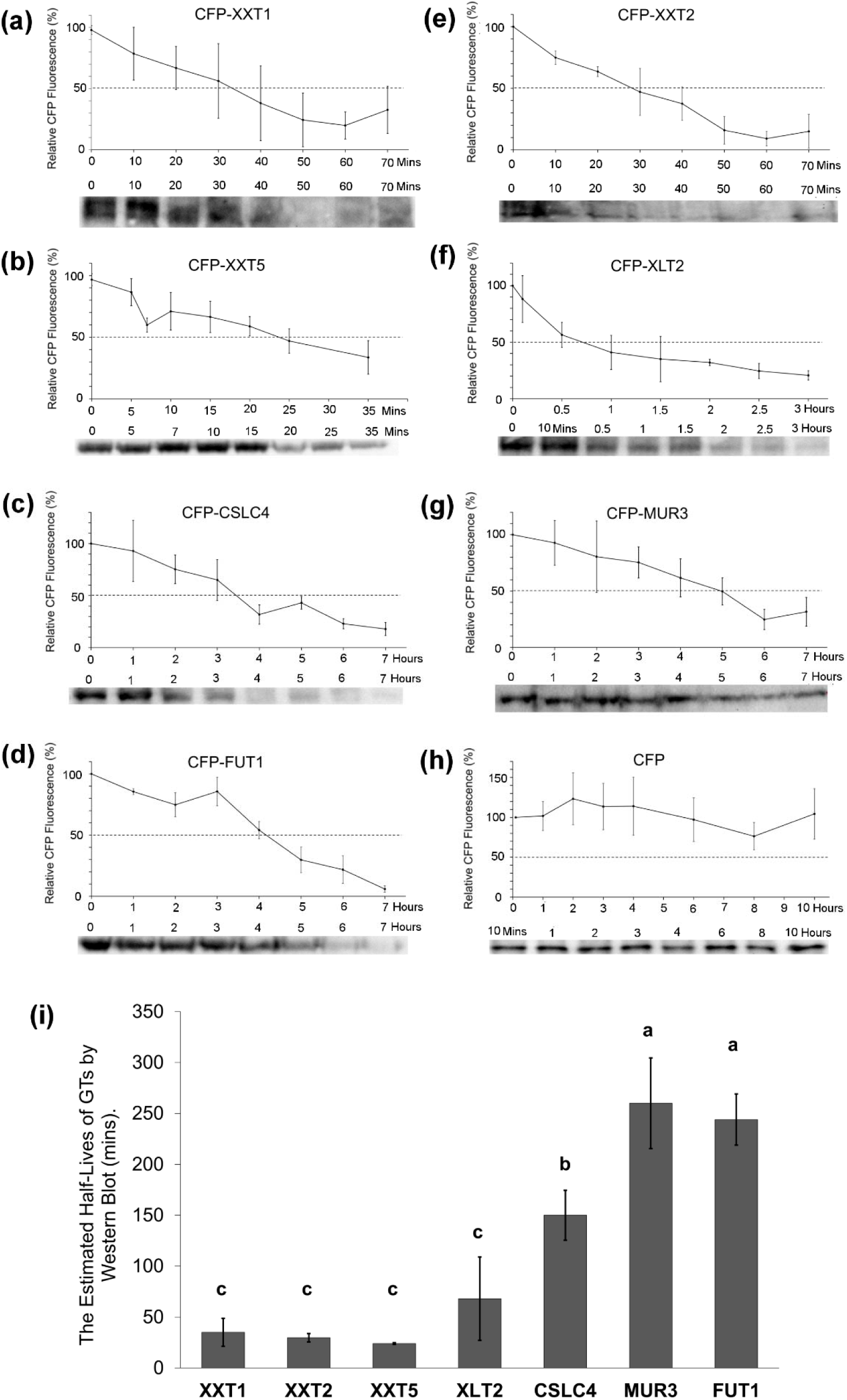
The estimation of the half-lives of GTs via Western Blot. The fourteen-day-old *Arabidopsis* seedlings were treated with 500 µg/mL CHX, and the CFP-GTs proteins were detected via Western Blot. The band intensity corresponding to the correct size of protein was analyzed by Image Fiji. The graphs show the intensity of bands on the membrane at different time points, and the data represents the means ±SD of three biological replicates (n=3). The dashed line indicated 50% reduction of band intensity. One representative example of Western Blot is shown under each graph for each time point. (a) The half-life of CFP-XXT1. (b) The half-life of CFP-XXT5. (c) The half-life of CFP-CSLC4. (d) The half-life of CFP-FUT1. (e) The half-life of CFP-XXT2. (f) The half-life of CFP-XLT2. (g) The half-life of CFP-MUR3. (h) The half-life of CFP. (i) The statistical analysis of the half-lives (T1/2) of GTs (n=3) using one-way ANOVA. Letter indicates the significant difference by one-way ANOVA (Tukey’s HSD p < 0.05).

### Estimation of the half-lives of XyG-synthesizing GTs using fluorescent microscopy

The measurement of the fluorescent signals from the CFP-GTs fusion proteins expressed in *Arabidopsis* KO mutant plants was used to estimate the half-lives of XyG-synthesizing enzymes in the root tips of seven-day-old seedlings. The intensity of the CFP signal in the root tips of the CFP-GT-expressing seedlings was measured at the time points specified in the figures during their continuous incubation in CHX-containing buffer (as described in the Materials and Methods section). The treated seedling roots were imaged until no detectable fluorescence was observed (Fig. 4). The 50% reduction in fluorescence intensity of XXT1 occurred around 30 mins (Fig. 4a), though some fluorescence was detected longer; it was reduced to the background level after a few hours. The half-life of XXT2 was 45 mins (Fig. 4e) and the complete degradation of XXT2 might require a longer time than XXT1 since the residual 10% fluorescence was observed after 8 h of incubation with CHX in comparison with XXT1. The half-life of XXT5 was about 20 mins, which is the shortest among all XyG-synthesizing proteins (Fig. 4b). The complete degradation of XXT5 protein was faster than XXT1 and XXT2, and almost no fluorescence signal of CFP-XXT5 was observed after 1 h of CHX treatment (Fig. 4b). The half-life of XLT2 was about 45 mins and somewhat longer than XXTs (Fig. 3f and 4f). A low CFP signal was detected after 4 h of incubation with CHX, and no CFP signal of GFP-XLT2 was detected after 8 h of treatment. For the long-living group, the half-life of CSLC4 was about 3.5 h, the half-life of FUT1 was about 4.0 h, and the half-life of MUR3 was about 5.5 h (Fig. 4c, d and g). The half-life of free CFP was much longer, and 90% of the CFP fluorescence signal remained after 12 h of incubation of the seedlings in the buffer containing CHX (Fig. 4h).

**Figure 4.**
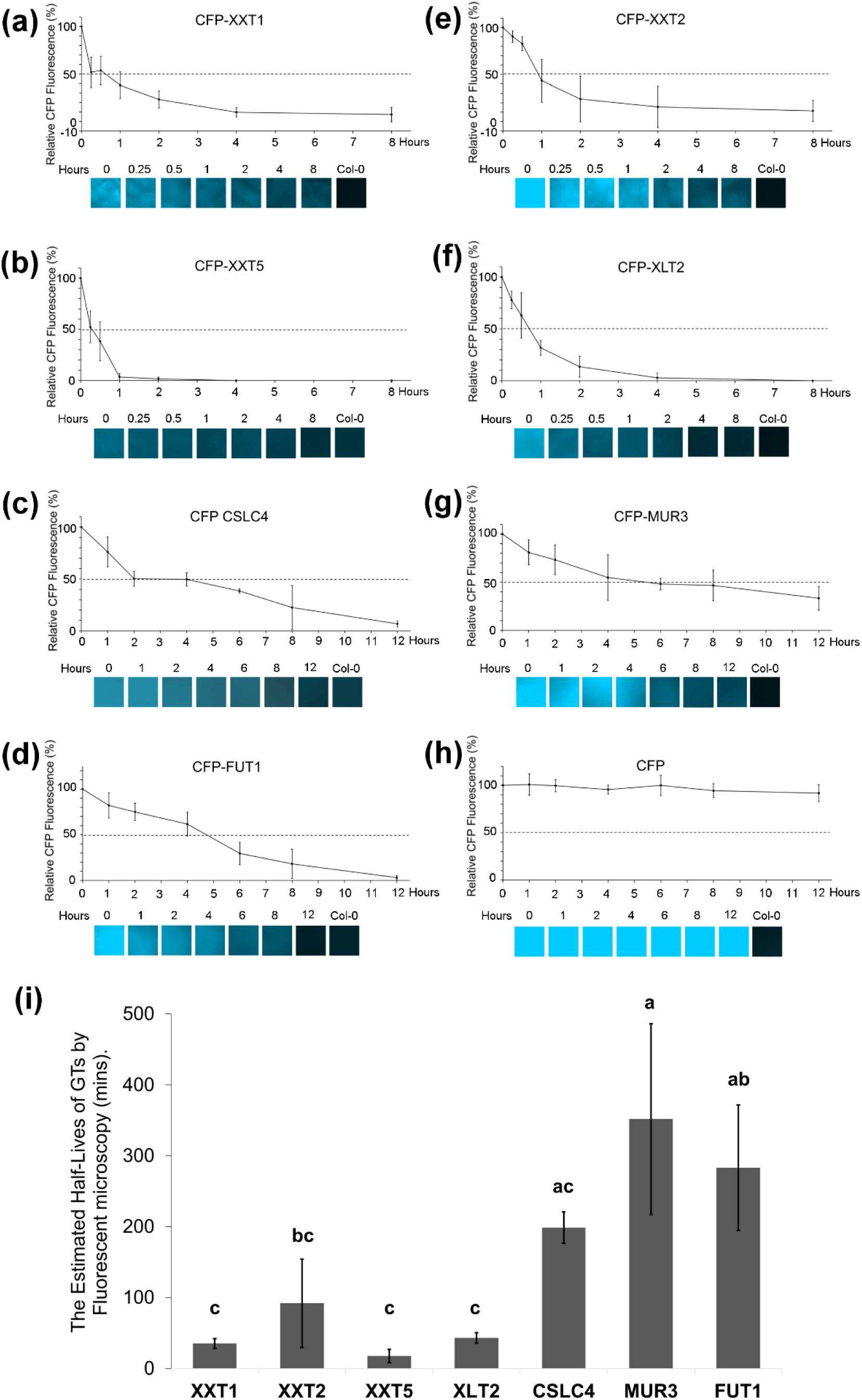
The estimation of the half-lives of GTs via fluorescent microscopy. The seven-day-old *Arabidopsis* seedlings were treated with 500 µg/mL CHX, and the CFP-GTs proteins were detected via fluorescent microscopy. Image Fiji analyzed the CFP fluorescence intensity of CFP-GTs. The graphs show the CFP fluorescence intensity at different time points, and the data represent the means ±SD of three biological replicates (n=3). The dashed line indicated 50% of CFP fluorescence intensity. One representative example of the microscope image is shown under each graph for each time point. (a) The half-life of CFP-XXT1. (b) The half-life of CFP-XXT5. (c) The half-life of CFP-CSLC4. (d) The half-life of CFP-FUT1. (e) The half-life of CFP-XXT2. (f) The half-life of CFP-XLT2. (g) The half-life of CFP-MUR3. (h) The half-life of CFP. (i) The statistical analysis of the half-lives (T1/2) of GTs via fluorescent microscope (n=3) via one-way ANOVA. Letter indicates the significant difference by one-way ANOVA (Tukey’s HSD p < 0.05).

## Discussion

As previously reported, GTs responsible for XyG synthesis localize to the Golgi apparatus, where their protein–protein interactions indicate assembly into multiprotein complexes. However, it is not known whether GTs involved in XyG biosynthesis require preformation of some complexes in ER, similar to what was reported for xylan-synthesizing proteins. In addition, we know nothing about the stability of polysaccharide-synthesizing GTs and the speed of their degradation. To fill these gaps, in this study, we investigated whether their protein-protein interactions are initiated before they are delivered to the Golgi; what their spatial distribution and localization within the Golgi, and how long these enzymes last before they are degraded. GTs reach the Golgi apparatus via independent anterograde transport, adopting specific sub-Golgi distributions that enable their assembly into multiprotein complexes within defined cisternae. A recent study demonstrated that two XXTs carry two-arginine motifs in their cytoplasmic N-termini, which are critical for their Golgi localization and interactions with one of the COPII complex members, SarI (Zhang et al., 2026). The protein positioning in the Golgi decides the interactions among different GTs and their dwell time in individual stacks, thereby dominating enzymatic performance. Protein stability is estimated as half-lives to control GTs’ persistence and degradation dynamics, affecting the assembly, maintenance, and disassembly of XyG-synthesising complexes. Overall, discrete trafficking routes, stack-specific localization, and turnover kinetics form an essential scaffold for the dynamic construction and operation of Golgi-based multiprotein assemblies.

First, we demonstrated that XyG-synthesizing GTs do not depend on their mutual protein-protein interactions to be transported to the Golgi, which suggests their heterocomplex formation is initiated only when proteins localize in the Golgi. We found that the lack of all five CSLC proteins did not affect the localization of other XyG-synthesizing GTs in the Golgi. Similarly, expression of all YFP/CFP-GTs in the protoplasts prepared from the different single T-DNA mutants of *XXT2*, *XXT5*, *MUR3*, *XLT2*, and *FUT1* showed their localization in the Golgi, and none of them were localized in the ER. However, we cannot exclude the possibility that other unknown proteins serve as scaffolds for the complex. It would be interesting to extend this study in the future to investigate such option. In this regard, the delivery mechanism of GTs involved in XyG biosynthesis differs from that of GTs involved in the synthesis of other major hemicellulose, xylan. Previous studies proposed that the putative xylan glycosyltransferase TaGT43-4 serves as the scaffold protein, interacting with the glycosyltransferase TaGT47-13, mutase TaGT75-3, mutase TaGT75-4, and protein TaVER2, collectively forming a heterocomplex within the ER and recruiting it to Golgi (Jiang et al., 2016). Similarly, xylan-synthesizing GTs (AoIRX9, AoIRX14A, and AoIRX10) in Asparagus formed the protein complex and AoIRX9 drives the protein complex to the Golgi (Zeng et al., 2016).

The distribution of several XyG-synthesizing GTs and the carbohydrate epitopes of XyG have been investigated earlier (Chevalier et al., 2010; Parsons et al., 2019), and the findings suggested that XyG synthesis occurs sequentially in discrete Golgi cisternae. Here, we extended such a localization study to all seven XyG-synthesizing enzymes and used two different Golgi markers combined with confocal microscopy. Our results further confirmed that seven GTs exhibit differential subcellular localization within the Golgi, which suggests the differential distribution of protein complexes responsible for the synthesis of different XyG side chains. Earlier, it was reported that the xylosylated forms of the XyG glucans accumulated mostly in the medial Golgi (Parsons et al., 2019), and the results of immunogold localization of XXT1 showed its localization in the cis-/medial Golgi (Chevalier et al., 2010). Localization of XXT2 and XXT5 was not investigated in that study. The functions of XXT1 and XXT2 are partially redundant (Cavalier et al., 2008), and our results demonstrated that XXT1 and XXT2 are predominantly localized in cis/medial Golgi (Fig. 2). The half-lives of these XXTs were about 30 mins (Fig. 3 and 4), indicating their comparably shorter-term presence in the Golgi. These results suggest that XXTs, involved in the early steps of XyG biosynthesis, initiate the branching of the XyG backbone in the cis-/medial Golgi and, most likely, leave the XyG complex in the later steps of synthesis. It is possible that after the xylosylation of the glucan backbone is accomplished, the XXTs are quickly degraded. The XXT5 protein has the shortest half-life among the XXTs studied here and XXT5 also shows mainly cis and medial-Golgi localization. Recently, it was confirmed that XXT5, together with XXT3 and XXT4, was responsible for the addition of a third xylosyl to complete the synthesis of XXXG-type XyG (Zhang et al., 2023). XXT5 requires the products of XXT1 and XXT2 enzymatic activity, the XXGG-type substitution of the glucan backbone. The slightly shorter longevity of XXT5 suggests either a shorter presence in the complex or faster degradation compared with XXT1 and XXT2, which are required to add the first two xylosyl residues to the glucan. It is known that there is no XyG in the absence of XXT1 and XXT2 (Cavalier et al., 2008).

The medium-size XyG epitopes, with xylosyl and galactosyl residues added, accumulated in the trans-Golgi (Parsons et al., 2019), and the localization of MUR3 (Chevalier et al., 2010) was shown primarily in medial/trans-Golgi (Fig. 2). Localizations of both XLT2 and MUR3 in the Golgi do not depend on protein-protein interactions with other GTs. The 45-minute half-life of XLT2 is longer than the half-life of XXTs, which correlates with the sequential processing of XyG biosynthesis. We propose that XLT2, as an enzyme involved in the addition of galactosyl residues after xylosylation of the glucan backbone, is present in the complex up to the medial/trans-Golgi.

The completely-branched fucosylated XyG epitopes have been shown to accumulate in the trans-Golgi (Parsons et al., 2019), and FUT1 is mostly localized in the trans-Golgi (Fig. 4) (Chevalier et al., 2010). From our study, FUT1 is a relatively longer-living GT with a half-life of about 4-5 h (Fig. 2d and 3d), indicating its longer presence in the Golgi. We showed that the transportation of FUT1 from the ER to the Golgi is independent of other enzymes but requires the product of their enzymatic actions in the Golgi. It is possible that FUT1 is recycled for a longer time within the medial/trans-Golgi space and continuously added to the complex, while MUR3 completes the galactosylation of XyG, which serves as an acceptor for FUT1 to link fucose at the last step of synthesis. An earlier study demonstrated that FUT1, MUR3, and CSLC4 have strong protein-protein interactions (Chou et al., 2015).

To our surprise, MUR3, the second galactosyltransferase responsible for adding the second galactosyl residue, showed the longest stability. At this time, it is difficult to explain the significant difference in stability between the two galactosyltransferases involved at the same stage of XyG decoration. There are several possible reasons for MUR3’s long stability. For example, MUR3 is a much slower enzyme and requires a longer time to complete the synthesis of XLLD-type XyG.Recent in vitro studies demonstrated the activities of XLT2 and MUR3. These studies showed that the two enzymes act independently to add galactose to their respective positions in xylosylated glucan substates, thereby synthesizing XLXG, XXLG, or XLLG products. MUR3 required a longer chain of xylosylated glucan compared with XLT2 (Corulli et al., 2026; Zhong et al., 2026). Therefore, MUR3 has to remain in the complex for a longer time. Other possible explanations are that the MUR3 may undergo recycling back to the medial Golgi for a longer period or that its degradation may take longer.

CSLC4 is an integral protein with multiple transmembrane domains. The distribution of CSLC4 in the Golgi cisternae was investigated in this study for the first time. Our results showed its mainly cis/medial-Golgi localization (Fig. 4). CSLC’s long half-life of 3.5 h (Fig. 2c and 3c) and multiple protein-protein interactions with other XyG-synthesizing GTs (Chou et al., 2012, 2015) suggest that being involved in the very first step of XyG biosynthesis to synthesize the glucan backbone, it most likely functions as a central/scaffold protein of the complex. We hypothesize that the synthesis of long polymeric XyG requires the processive synthesis of a glucan backbone that is simultaneously branched by the action of other GTs, likely organized around CSLC4. The long life of CSLC4 supports the notion that it acts within cis-Golgi and medial Golgi, whereas, in the trans-Golgi, MUR3 and FUT1 might act within a smaller heterocomplex to complete the synthesis.

We compared the expression of all XyG-synthesizing GTs using data from the *Arabidopsis* RNA-seq database (http://ipf.sustech.edu.cn/pub/athrdb/) and the datasets for whole plants. Taking data from three replication libraries (DRX078165, DRX078166, and DRX078167), we estimated that the levels of expression were as follows: CSLC4 (71.6 FPKM), XXT2 (67.8 FPKM), and XXT5 (77.8 FPKM) were highly expressed, and the expression of XXT1 was slightly lower (40.3 FPKM). The expression levels of XLT2 (22.9 FPKM), MUR3 (31.7 FPKM), and FUT1 (20.1 FPKM) were the lowest. It is possible that the high levels of XXT proteins and their relatively higher activities enable rapid xylosylation of the glucan backbone in the earlier cisternae (cis/medial), after which the proteins can be degraded. The relatively lower expression of MUR3 and FUT1 may necessitate their longer residence times and recycling to complete the synthesis of fully branched XyG molecules before they are delivered to the cell surface.

Although the mechanism of stepwise synthesis and branching of XyG can be considered as one plausible explanation for the observed differences in GTs’ half-lives, their different recycling patterns may be another contributing factor. Currently, we do not have any information about the recycling mechanisms of GTs localized in the Golgi. According to the accepted notion, the plant Golgi undergoes the maturation process of its stacks from the cis to medial to trans-Golgi (Ito et al., 2014; Kurokawa et al., 2019; Robinson, 2020). Due to this maturation mode, the cis-cisternae content, including the stable protein complexes of GTs, can move into the medial- and trans-cisternae. This Golgi maturation process can support the differential distribution of XyGs bearing different branching patterns, as well as the differential distribution of the enzymes that synthesize them. Thus, the potentially different GT recycling mechanisms may also affect their half-lives.

Here, we demonstrated that GTs involved in XyG biosynthesis are independently transported to the Golgi apparatus, where they reside for different durations in distinct Golgi cisternae and, together with other interacting proteins, participate in XyG biosynthesis. Currently, we cannot conclusively propose all contributing factors that determine these reorganizations, but it is plausible to suggest that the XyG synthesis follows the stepwise process of decorating the glucan backbone with longer side chains in a manner similar to that proposed for the processes of protein glycosylation (Schoberer et al., 2013; Strasser et al., 2021, Preprint; Zabotina et al., 2021). This study advances our mechanistic understanding of XyG biosynthesis, specifically shedding light on the functional organization and degradation of the major enzymes involved in this process within the plant Golgi. Uncovering more details about polysaccharide biosynthesis will enable more informed modifications of these processes to generate new compositions of cell wall polysaccharides and improve plant fitness and biomass application for renewable materials.

## Abbreviations

AoIRX: Irregular xylem CFP
Cyan: Fluorescent Protein
CHX: Cycloheximide
CSLC: Cellulose synthase-like
GTs: Glycosyltransferases
KO: Knockout
SD: Standard deviations
UGTs: UDP-glucuronosyltransferases
XXT: Xyloglucan xylosyltransferases
XyGs: Xyloglucans
YFP: Yellow fluorescent protein

## Short legends for Supporting Information

Supporting Table. S1 The primers used in the study.

Supporting Figure. S1 Transient co-expression of CFP/YFP-GTs with Golgi marker Golgi-Y (YFP) / R (mCherry) in *cslc456812 Arabidopsis* protoplasts. Bar, 10 μm.

Supporting Figure. S2 Transient co-expression of CFP/YFP-GTs with Golgi marker Golgi-Y (YFP) / R (mCherry) in *xlt2 Arabidopsis* protoplasts. Bar, 10 μm.

Supporting Figure. S3 Transient co-expression of CFP/YFP-GTs and Golgi marker Golgi-R(mCherry) in *mur3 Arabidopsis* protoplasts. Bar, 10 μm.

Supporting Figure. S4 Transient co-expression of CFP/YFP-GTs with Golgi marker Golgi-Y (YFP) / R (mCherry) in *fut1 Arabidopsis* protoplasts. Bar, 10 μm.

Supporting Figure. S5 Transient co-expression of CFP/YFP-GTs with Golgi marker Golgi-Y (YFP) / R (mCherry) in *xxt2 Arabidopsis* protoplasts. Bar, 10 μm.

Supporting Figure. S6 The localization analysis of YFP-XXT2 with cis-Golgi marker Golgi-R.

Supporting Figure. S7 The localization analysis of YFP-XXT5 with cis-Golgi marker Golgi-R.

Supporting Figure. S8 The localization analysis of YFP-CSLC4 with cis-Golgi marker Golgi-R.

Supporting Figure. S9 The localization analysis of YFP-XLT2 with cis-Golgi marker Golgi-R.

Supporting Figure. S10 The localization analysis of YFP-MUR3 with cis-Golgi marker Golgi-R.

## Acknowledgments

We thank Margaret Carter for her technical assistance with fluorescent microscopy.

## Author contributions

OAZ conceived the research ideas and developed the project. NZ and KU did the experiments and collected the data. NZ wrote the manuscript. OAZ, and NZ contributed to manuscript preparation.

## References

Anderson CT. 2016. We be jammin’: An update on pectin biosynthesis, trafficking and dynamics. Journal of experimental botany 67, 495–502.

Averbeck N, Gao X-D, Nishimura S-I, Dean N. 2008. Alg13p, the Catalytic Subunit of the Endoplasmic Reticulum UDP-GlcNAc Glycosyltransferase, Is a Target for Proteasomal Degradation. Molecular Biology of the Cell 19, 2169–2178.

Barbier O, Be A, Langer, Hum DW. 1999. Cloning and characterization of a simian UDP-glucuronosyltransferase enzyme UGT2B20, a novel C 19 steroid-conjugating protein. The Biochemical journal 337, 567–574.

Barbier O, Girard C, Breton R, et al. 2000. N-glycosylation and residue 96 are involved in the functional properties of UDP-glucuronosyltransferase enzymes. Biochemistry 39, 11540–11552.

Cavalier DM, Lerouxel O, Neumetzler L, et al. 2008. Disrupting two Arabidopsis thaliana xylosyltransferase genes results in plants deficient in xyloglucan, a major primary cell wall component. Plant Cell 20,1519–1537.

Chevalier L, Bernard S, Ramdani Y, et al. 2010. Subcompartment localization of the side chain xyloglucan-synthesizing enzymes within Golgi stacks of tobacco suspension-cultured cells. Plant Journal 64, 977–989.

Chou YH, Pogorelko G, Young ZT, et al. 2015. Protein-protein interactions among xyloglucan-synthesizing enzymes and formation of golgi-localized multiprotein complexes. Plant and Cell Physiology 56, 255–267.

Chou YH, Pogorelko G, Zabotina OA. 2012. Xyloglucan xylosyltransferases XXT1, XXT2, and XXT5 and the glucan synthase CSLC4 form Golgi-localized multiprotein complexes. Plant Physiology 159. 1355–1366.

Clough SJ, Bent AF. 1998. Floral dip: A simplified method for Agrobacterium-mediated transformation of Arabidopsis thaliana. Plant Journal 16. 735–743.

Cocuron J-C, Lerouxel O, Drakakaki G, et al. 2007. A gene from the cellulose synthase-like C family encodes a-1,4 glucan synthase. Proceedings of the National Academy of Sciences of the United States of America 104, 8550–8555.

Corulli CJ, Graf AS, Chapla D, et al. 2026. Biochemical characterization of xyloglucan galactosyltransferases MUR3 and XLT2 from Spirodela polyrhiza. The Plant journal: for cell and molecular biology. 125, e70754.

Gillespiesq W, Kelmsil S, Paulson JC. 1992. Cloning and expression of the Gal beta 1, 3GalNAc alpha 2,3-sialyltransferase. The Journal of biological chemistry 267, 21004–21010.

Grefen C, Donald N, Hashimoto K, et al. 2010. A ubiquitin-10 promoter-based vector set for fluorescent protein tagging facilitates temporal stability and native protein distribution in transient and stable expression studies. The Plant journal: for cell and molecular biology 64, 355–365.

Gupta R, Leon F, Rauth S, et al. 2020. A Systematic Review on the Implications of O-linked Glycan Branching and Truncating Enzymes on Cancer Progression and Metastasis. Cells 9, 446.

Ito Y, Uemura T, Nakano A. 2014. Formation and maintenance of the golgi apparatus in plant cells. International Review of Cell and Molecular Biology. Elsevier Inc., 221–287.

Jensen JK, Schultink A, Keegstra K, et al. 2012. RNA-seq analysis of developing nasturtium seeds (Tropaeolum majus): Identification and characterization of an additional galactosyltransferase involved in xyloglucan biosynthesis. Molecular plant 5, 984–992.

Jiang N, Wiemels RE, Soya A, et al. 2016. Composition, assembly, and trafficking of a wheat xylan synthase complex. Plant Physiology 170, 1999–2023.

Julian JD, Zabotina OA. 2022. Xyloglucan Biosynthesis: From Genes to Proteins and Their Functions. Frontiers in Plant Science 13, 920494.

Kim S-J, Chandrasekar B, Rea AC, et al. 2020. The synthesis of xyloglucan, an abundant plant cell wall polysaccharide, requires CSLC function. Proceedings of the National Academy of Sciences of the United States of America. 117, 20316–20324.

Kitano M, Kizuka Y, Sobajima T, et al. 2021. Rab11-mediated post-Golgi transport of the sialyltransferase ST3GAL4 suggests a new mechanism for regulating glycosylation. Journal of Biological Chemistry 296. 100354.

Kong Y, Peña MJ, Renna L, et al. 2015. Galactose-depleted xyloglucan is dysfunctional and leads to dwarfism in arabidopsis. Plant Physiology 167. 1296–1306.

Kurokawa K, Osakada H, Kojidani T, et al. 2019. Visualization of secretory cargo transport within the Golgi apparatus. Journal of Cell Biology 218, 1602–1618.

Li Q, Ran P, Zhang X, et al. 2018. Downregulation of N-Acetylglucosaminyltransferase GCNT3 by miR-302b-3p Decreases Non-Small Cell Lung Cancer (NSCLC) Cell Proliferation, Migration and Invasion. Cellular Physiology and Biochemistry 50, 1005–1014.

Lund CH, Bromley JR, Stenbæk A, et al. 2015. A reversible Renilla luciferase protein complementation assay for rapid identification of protein-protein interactions reveals the existence of an interaction network involved in xyloglucan biosynthesis in the plant Golgi apparatus. Journal of Experimental Botany 66, 85–97.

Madson M, Dunand C, Li X, et al. 2003. The MUR3 gene of Arabidopsis encodes a xyloglucan galactosyltransferase that is evolutionarily related to animal exostosins. Plant Cell 15, 1662–1670.

Myoung Shin E, Thang Huynh V, Abda Neja S, et al. 2021. GREB1: An evolutionarily conserved protein with a glycosyltransferase domain links ER glycosylation and stability to cancer. Science advances 7, eabe2470.

Nelson BK, Cai X, Nebenführ A. 2007. A multicolored set of in vivo organelle markers for co-localization studies in Arabidopsis and other plants. Plant Journal 51, 1126–1136.

Parsons HT, Stevens TJ, McFarlane HE, et al. 2019. Separating Golgi proteins from cis to trans reveals underlying properties of cisternal localization. Plant Cell 31, 2010–2034.

Pauly M, Keegstra K. 2016. Biosynthesis of the Plant Cell Wall Matrix Polysaccharide Xyloglucan. Annual review of plant biology 67, 235–259.

Peña MJ, Ryden P, Madson M, et al. 2004. The Galactose Residues of Xyloglucan Are Essential to Maintain Mechanical Strength of the Primary Cell Walls in Arabidopsis during Growth. Plant Physiology 134, 443–451.

Peng K, Liu R, Jia C, et al. 2021. Regulation of o-linked n-acetyl glucosamine transferase (Ogt) through e6 stimulation of the ubiquitin ligase activity of e6ap. International Journal of Molecular Sciences 22, 10286.

Robinson DG. 2020. Plant Golgi ultrastructure. Journal of Microscopy 280, 111–121.

Scheller HV, Ulvskov P. 2010. Hemicelluloses. Annual Review of Plant Biology 61, 263–289.

Schoberer J, Liebminger E, Botchway SW, et al. 2013. Time-resolved fluorescence imaging reveals differential interactions of N-glycan processing enzymes across the golgi stack in planta. Plant Physiology 161, 1737–1754.

Strasser R, Seifert G, Doblin MS, et al. 2021. Cracking the “Sugar Code”: A Snapshot of N- and O-Glycosylation Pathways and Functions in Plants Cells. Frontiers in Plant Science 12, 640919.

Sumardika IW, Youyi C, Kondo E, et al. 2018. β-1,3-galactosyl-o-glycosyl-glycoprotein β-1,6-N-acetylglucosaminyltransferase 3 increases MCAM stability, which enhances S100A8/A9-mediated cancer motility. Oncology Research 26, 431–444.

Tamura K, Shimada T, Kondo M, et al. 2005. KATAMARI1/MURUS3 is a novel Golgi membrane protein that is required for endomembrane organization in Arabidopsis. Plant Cell 17, 1764–1776.

Turgeon D, Bastien Carrier J-S, Lé Vesque R, et al. 2001. Relative Enzymatic Activity, Protein Stability, and Tissue Distribution of Human Steroid-Metabolizing UGT2B Subfamily Members. Endocrinology 142, 778–787.

Vaidyanathan K, Niranjan T, Selvan N, et al. 2017. Identification and characterization of a missense mutation in the O-linked β-N-acetylglucosamine (O-GlcNAc) transferase gene that segregates with X-linked intellectual disability. Journal of Biological Chemistry 292, 8948–8963.

Vanzin GF, Madson M, Carpita NC, et al. 2002. The mur2 mutant of Arabidopsis thaliana lacks fucosylated xyloglucan because of a lesion in fucosyltransferase AtFUT1. Proceedings of the National Academy of Sciences of the United States of America 99, 3340–3345.

Vuttipongchaikij S, Brocklehurst D, Steele-King C, et al. 2012. Arabidopsis GT34 family contains five xyloglucan α-1,6-xylosyltransferases. New Phytologist 195, 585–595.

Zabotina OA, Avci U, Cavalier D, et al. 2012. Mutations in multiple XXT genes of arabidopsis reveal the complexity of xyloglucan biosynthesis. Plant Physiology 159, 1367–1384.

Zabotina OA, Van De Ven WTG, Freshour G, et al. 2008. Arabidopsis XXT5 gene encodes a putative α-1,6-xylosyltransferase that is involved in xyloglucan biosynthesis. The Plant journal: for cell and molecular biology 56, 101–115.

Zabotina OA, Zhang N, Weerts R. 2021. Polysaccharide Biosynthesis: Glycosyltransferases and Their Complexes. Frontiers in Plant Science 12, 625307.

Zeng W, Lampugnani ER, Picard KL, et al. 2016. Asparagus IRX9, IRX10, and IRX14A are components of an active xylan backbone synthase complex that forms in the Golgi apparatus. Plant Physiology 171, 93–109.

Zhang N, Julian JD, Yap CE, et al. 2023. The Arabidopsis xylosyltransferases, XXT3, XXT4, and XXT5, are essential to complete the fully xylosylated glucan backbone XXXG-type structure of xyloglucans. The New phytologist 238, 1986–1999.

Zhang N, Julian JD, Zabotina OA. 2026. Interaction with COPII Member SAR1 Is Critical for the Delivery of Arabidopsis Xyloglucan Xylosyltransferases XXT2 and XXT5 to the Golgi Apparatus. Plants 15, 822.

Zhao T, Zhao X, Qian K, et al. 2022. Radiotherapy prognosis-associated gene GCNT3 promotes the proliferation, migration and invasion of lung adenocarcinoma cells. Heliyon 8, e12100.

Zhong R, Phillips DR, Ye Z-H. 2026. Biochemical insights into the regiospecificity of xyloglucan galactosyltransferases. (S Turner, Ed.). Journal of Experimental Botany erag117. Advance online publication.

